# Multiset correlation and factor analysis enables exploration of multi-omic data

**DOI:** 10.1101/2022.07.18.500246

**Authors:** Brielin C. Brown, Collin Wang, Silva Kasela, François Aguet, Daniel C. Nachun, Kent D. Taylor, Russell P. Tracy, Peter Durda, Yongmei Liu, W. Craig Johnson, David Van Den Berg, Namrata Gupta, Stacy Gabriel, Joshua D. Smith, Robert Gerzsten, Clary Clish, Quenna Wong, George Papanicolau, Thomas W. Blackwell, Jerome I. Rotter, Stephen S. Rich, Kristin G. Ardlie, David A. Knowles, Tuuli Lappalainen

## Abstract

Multi-omics datasets are becoming more common, necessitating better integration methods to realize their revolutionary potential. Here, we introduce Multi-set Correlation and Factor Analysis, an unsupervised integration method that enables fast inference of shared and private factors in multi-modal data. Applied to 614 ancestry-diverse participant samples across five ‘omics types, MCFA infers a shared space that captures clinically relevant molecular processes.

## Main

Recent years have seen an explosion in multi-omic data, with studies simultaneously profiling RNA expression, protein levels, chromatin accessibility and more^1^. By providing complementary views into the underlying biology, these datasets promise to illuminate molecular processes and disease states that cannot be gleaned from any lone modality^2^. However, joint inference methods are lacking in either the number or type of modes that can be used, or in flexibility and efficiency^1^. Multi-omic data bring substantial challenges: distributions differ between modes, the sample size is typically small relative to features, efficient algorithms are needed, and each mode has contributions from factors that are shared between modes and unique to itself^3,4^. Canonical correlation analysis (CCA) is a statistical technique that infers shared factors between two data modes by finding correlated linear combinations of the features in each^5^. CCA has enjoyed substantial attention in genomics^6–9^, however, extending CCA to additional modes is fraught; at least 10 different formulations are equivalent in the two-mode case^10^, and many are challenging to fit^11^. Equivalently, CCA can be conceptualized as a probabilistic model (pCCA), revealing a connection to factor analysis^12^.

We have developed Multiset Correlation and Factor Analysis (MCFA, Figure 1a), an unsupervised integration method that generalizes pCCA and factor analysis, enabling fast inference of shared and private factors in multimodal data. MCFA is based on two insights: 1) unlike traditional CCA, pCCA has only one natural extension to multi-modal data, which is both conceptually elegant and efficient to fit, and 2) after fitting pCCA, the residual in a mode represents private structure, which is well-modeled by factor analysis. Our method combines these insights to fit factors that are shared across modalities and private to each simultaneously. For efficiency and regularization, MCFA uses the top principal components (PCs) of each mode^6,7^. It allows use of random matrix techniques^13^ to choose the shared dimensionality and number of PCs, eliminating tuning parameters. Finally, MCFA is a natural approach to integration: in the supplemental note, we detail a theoretical connection between our model and multiset CCA.

**Figure 1:**
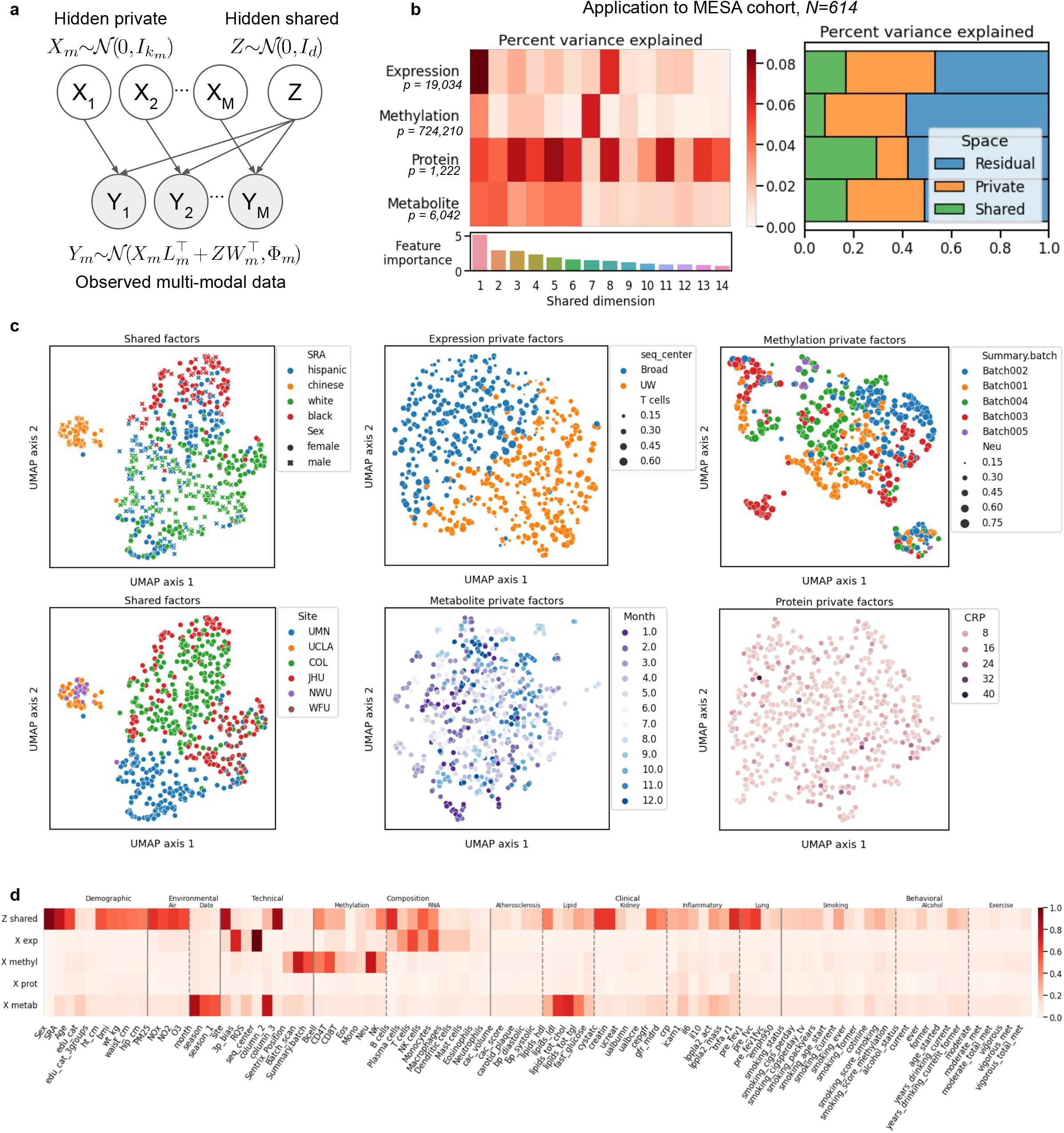
a) The MCFA model. b) Breakdown of the variance in four omics types captured by the inferred space. c) UMAP embedding of the shared and private spaces, annotated with the most relevant feature set. d) Variance in sample metadata explained by each learned space.

We have applied MCFA to 614 ancestry-diverse individuals from the Multi-Ethnic Study of Atherosclerosis (MESA)^14^, which has collected comprehensive phenotypic data of its subjects. The Trans-Omics for Precision Medicine (TOPMed)^15^ program instituted a multi-omics pilot study to evaluate the utility of long-term stored samples for discovery related to heart, lung, blood, and sleep disorders. MESA provided samples for five ‘omics types: 1) whole genome sequencing (WGS), 2) RNA-sequencing of peripheral blood mononuclear cells (PBMCs), 3) DNA methylation array profiling from whole blood 4) protein mass spectrometry of blood plasma, and 5) metabolite mass spectrometry of blood plasma. We integrated RNA-sequencing, methylation, protein, and metabolite data from Exam 1 using MCFA, which inferred a fourteen-dimensional shared space. We found that shared structure explained a large proportion of the variance in each mode (Figure 1b, right). Protein levels had the highest sharing with 29.2% of the variance explained (VE) by the shared space, followed by RNA and metabolite levels (16.6% and 17.1%, respectively). Methylation showed the least sharing, with only 8.1% VE by the shared space. Due to the high dimensionality of the data and the limited sample size, about half of the variance in each dataset is unmodeled to reduce overfitting. Using MCFA, it is possible to further infer the variance in each modality explained by the individual factors, thus determining which modalities contribute to each (Figure 1b, left). Our top factor has contributions from all modalities, but their respective contributions to the other factors vary substantially.

We used uniform manifold approximation and projection (UMAP)^16^ to construct a 2D embedding of the shared and private spaces (Figure 1c). We noticed a striking clustering of the individuals by self-reported ancestry (SRA) and sex in the shared space, even though the top PCs of individual modes do not cluster by these factors (Figure S1), and the shared space was inferred without genetic or sex chromosome features. Shared factor 1 separates Black and white individuals, with Hispanic individuals in between, while factor 3 separates Chinese individuals, and factor 2 differentiates by sex (Figure S1, S2). We validated this structure via leave-one-out cross-validation, indicating our PC selection strategy mitigated over-fitting (Figure S3).

Next, we evaluated the total phenotypic variance explained by each of our inferred spaces (Figure 1d, S2, Table S1-S3). The shared space captured 95.3% of the variation in sex, 83.3% in site, 80.0% in SRA, and 60.2% in age. The shared space also captured anthropomorphic differences such as BMI (51.0% VE) and clinical measures including those related to kidney function (creatine, 64.8% VE) and inflammation (TNF-alpha receptor-1 69.1% VE). We used CIBERSORT^17^ and the Houseman method^18^ to estimate the cell-type composition of our RNA (PBMC) and methylation (whole blood) samples, respectively. Both shared and privates spaces contributed to the relative proportions of PBMC-abundent cell types (e.g. T cells and NK cells) estimated from both data modalities, while the proportion of PBMC-depleted types (e.g. neutrophils) estimated from the methylation data was only captured by the methylation private space. Modality-private spaces frequently captured technical factors: 100% of the variance in sequencing center and 71.6% of the variance in 3-prime bias are captured by the RNA private space, while 76.8% of methylation array batch is captured by its private space. Many phenotypes that are themselves measurements of metabolites were captured by the metabolite private space, however the strongest association was with the month of sample collection (85.8% VE). We noticed no large associations between the protein private space and any of our metadata, despite several of our phenotypes being clinical protein markers, however, several of these factors are partially captured by the shared space.

Finally, we integrated WGS data by conducting a GWAS of the inferred factors while controlling for site, age, sex and 11 genotype PCs. Given our limited sample size, we did not expect to find genome-wide significant associations. However, we hypothesized that genetic associations with our inferred factors, which represent major axes of molecular variation, may be enriched for known GWAS hits or *trans*-eQTLs. We obtained a list of 10,174 such associations from the eQTLgen consortium^19^, of which 3,854 are trans-eQTLs, and further defined a more limited set of 1,107 “influential” *trans*-eQTLs that affect at least 10 genes. We tested the GWAS of each factor for enrichment of these three categories and found 9 significant enrichments (mean 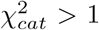, FDR 5%, Figure 2a, Figure S4). Factor 7 showed the strongest enrichment for reported GWAS hits and *trans*-eQTLs. The top GWAS SNPs associated with factor 7 are from blood lipid studies and are located primarily around the FADS1 and FADS2 genes that are known to regulate lipid metabolism^20^. These include rs174541 (*p* = 4.3 × 10^−5^ for factor 7 association) which is also reported in GWAS of type-2 diabetes^21^, rs174549 (*p* = 5.6 × 10^−5^) which is also reported in GWAS of white blood cell count^22^, and rs1535 (*p* = 8.3×10^−5^), which is also reported in GWAS of inflammatory bowel disease^23^ (Table S4). Factor 7 explains 6.7% of the modeled variation in methylation, the largest of any factor, and many of the top-associated SNPs were also cis-methylation QTLs in MESA^24^ (Figure 2c). Factor 7 is anti-correlated with sample proportion of CD8 T cells and NK cells estimated from methylation data (*ρ* = −0.41 and *ρ* = −0.25), and correlated with BMI (*ρ* = 0.25) and measures of inflammation including TNF-R1 (*ρ* = 0.33) and interleukin-6 (*ρ* = 0.20) (Figure 2b). While further research is needed to establish causal relationships of these genetic effects on methylation in cis and trans as well as on diverse traits, we note that DNA methylation patterns have been previously associated with lipid metabolism and metabolic disease^25,26^.

**Figure 2:**
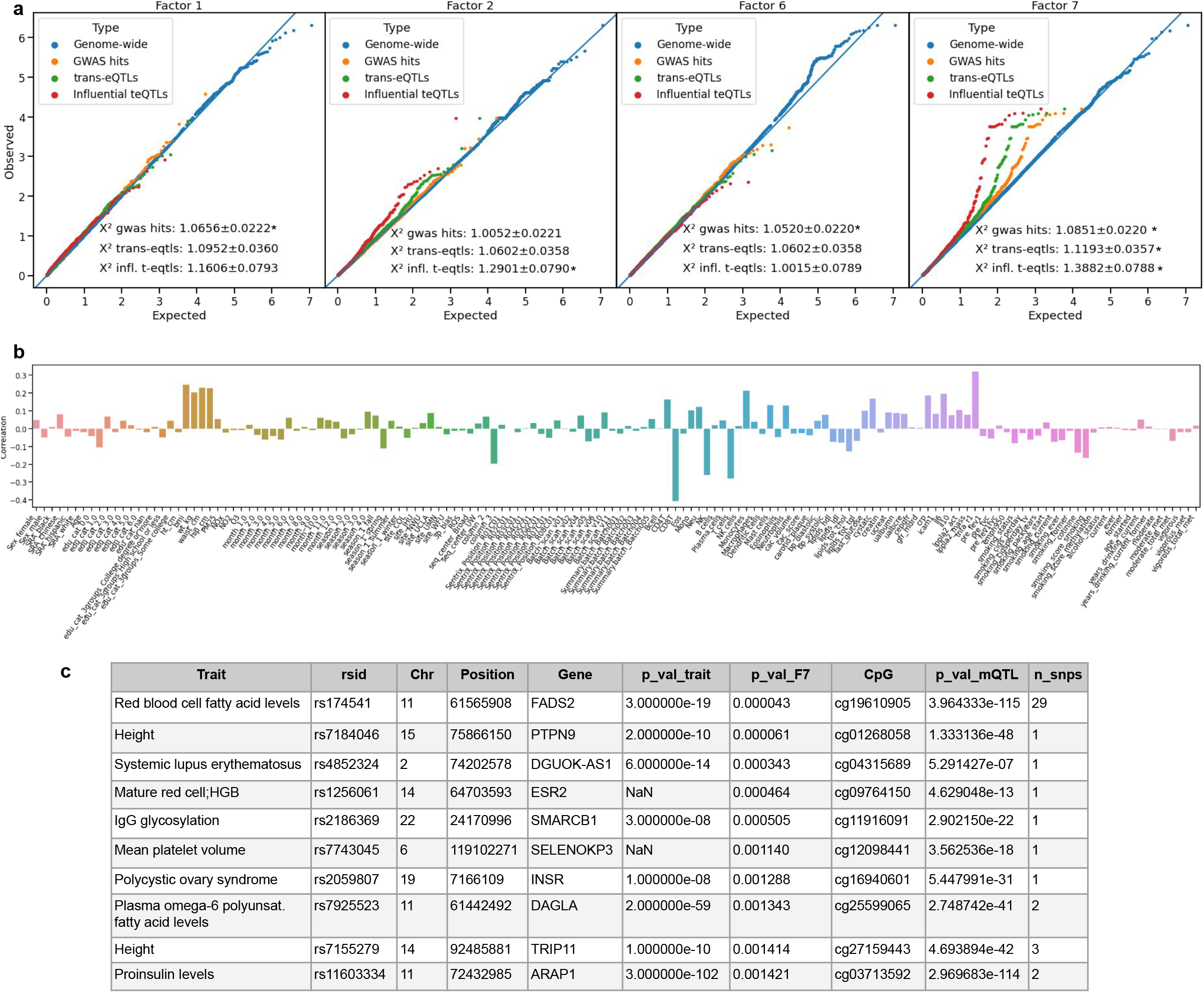
a) QQ-plot of a GWAS for factors 1, 2, 6, and 7. b) Correlation of factor 7 with sample metadata. c) Top unique SNP-CpG pairs for known trait-associated SNPs additionally associated with factor 7. `n_snps` indicates the number of SNPs suggestively-associated with factor 7 that have the same top-associated CpG.

MCFA has several advantages compared to other multi-omic integration approaches. Compared to group factor analysis methods^4^, MCFA separates modality-specific from dataset-shared factors. Compared to non-negative matrix factorization-based methods^3^ that share a feature weight set across modalities, MCFA is able to use all data types. Due to the use of observational data and unsupervised methods, all analyses should be considered exploratory; they can find structure in the data while generating hypotheses but cannot be used to make causal claims and may reflect properties of the underlying data. For example, in MESA the sample collection site is strongly correlated with self-reported ancestry (SRA) and air-quality. We repeated our analysis of the variance explained by the learned space while additionally controlling for site (Table S3), and noticed a small decrease in the proportion of VE in SRA (from 80.0% to 71.6%) and a large decrease in the variance explained in PM25 (from 66.8% to 24.2%). In this study air quality and site are nearly co-linear and thus their independent effects cannot be distinguished. Future work with larger sample sizes may allow for network inference methods to generate directed hypotheses^27^. Genetic associations are particularly valuable in this, with the inferred axes of molecular variation providing a promising future trait for GWAS and pheWAS studies. TOPMed is among the most ambitious current efforts to collect multi-omic population-level data, thus given the results of this pilot analysis we expect future integration studies in this cohort to be fruitful.

## 1 Online methods

### 1.1 Multiset correlation and factor analysis

Let 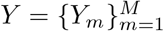 be a set of *N* × *p*_*m*_ observed data matrices: *N* individuals measured in *M* data modalities consisting of *p*_*m*_ features each. We model each observed mode as having contributions from two low-dimensional hidden factors (Figure 1a, Figure S7)

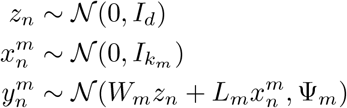

where *d* is the shared hidden dimensionality, *k*_*m*_ are the dataset-private hidden dimensionalities, *W*_*m*_ are *p*_*m*_ ×*d* shared space loading matrices, *L*_*m*_ are *p*_*m*_ × *k*_*m*_ private space loading matrices and 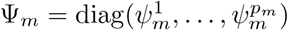 are the diagonal residual covariance matrices. Given *Y, d* and *k*_*m*_, our goal is to infer the hidden factors *Z* and *X*_*m*_ and loading matrices *W*_*m*_ and *L*_*m*_. This can be accomplished using a straightforward application of expectation-maximization (EM)^1^. For a derivation of the EM update equations, as well as a more detailed exposition including the relationship to pCCA, factor analysis and other multiset CCA (MCCA) methods, see the Supplemental Note. In practice, we center and scale all data variables. This is not strictly required, however it enables simple estimation of the number of PCs to include and simplifies explained variance calculations, see below.

### 1.2 Model initialization

An important aspect of EM optimization is choosing a good initialization. We benchmarked three approaches to initializing *W*: random initialization and two versions of MCCA that correspond to maximizing the sum of pairwise correlations with the average variance and average norm constraints. These MCCA fomulations can be solved via simple eigendecompositions. We found that the sum of pairwise correlations with average variance constraint produced the best initial estimates (Figure S5). This can be solved with a simple two step procedure: 1) whiten each data matrix using the singular value decomposition (SVD), 2) perform a second SVD on the concatenated whitened data matrices^2^:

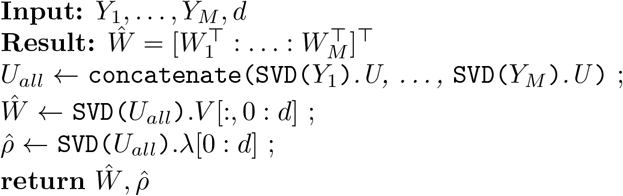

We initialize *L* and Ψ using probabilistic PCA on the residual data matrices after fitting MCCA. Specifically:

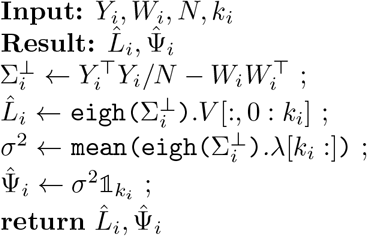

### 1.3 High dimensionality and selection of hyperparameters

There are two primary approaches to control for over-fitting in applications of CCA-type methods to high-dimensional (*N* ≪ *p*) problems. The first is to use penalized optimization techniques, where the objective function additionally contains an *l*_1_ constraint on the weight matrices^3^. The second is to project each dataset onto its *informative* principal components^4–6^. In this application, we choose the latter approach in order to find components with broad effects on the structure of the data, rather than specific effects on small numbers of molecular features^6^. We choose the number of principal components of each dataset using the Marchenko-Pasteur law^7^, which states that for mean 0, variance 1 data, principal components with corresponding eigenvalues above 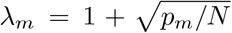 should be considered non-noise. We are not aware of a corresponding law for the cross-covariance matrices used in CCA, however, the empirical spectral distribution of the cross-covariance of matrices of random noise can be easily estimated in practice:

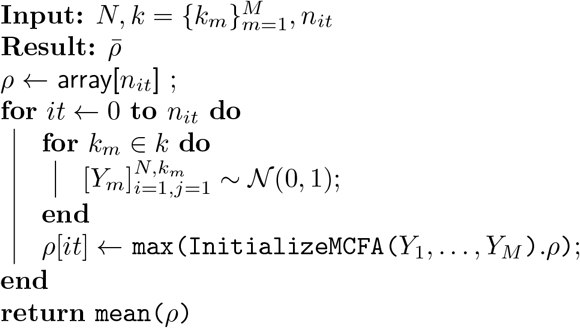

Then we keep all components where 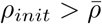.

### 1.4 Calculating the variance explained

The linear-Gaussian nature of the model simplifies estimation of the variance explained. That is, if the features of each mode 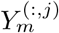 are normalized to variance 1, the model 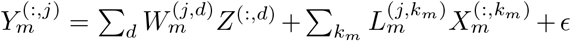 implies that the variance in feature *j* of mode *m* explained by shared factor *d* is 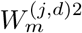. Likewise, the variance explained by the *k*_*m*_-th private factor of mode *m* is 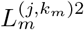. The total variance in mode *m* explained by a given shared factor *d* (respectively, private factor *k*_*m*_) is thus given by 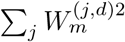 (respectively,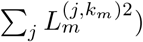), and the total variance in the mode explained by the factors are 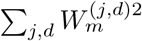 and 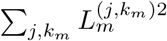, respectively. Note that when working in PC-space, the raw *W* and *L* features correspond to variance in PCs explained, rather than modality features. Thus, we calculate the variance explained after projecting back into the original feature space *W*_*m*_ ← *V*_*m*_*W*_*m*_, *L*_*m*_ ← *V*_*m*_*L*_*m*_ where *V*_*m*_ are the right singular vectors of mode *m*. To calculate the variance in a metadata feature explained by a particular space, we regressed the trait value *T* on the shared or private space, *T* ∼ *Z* or *T* ∼ *X*_*m*_. For continuous-valued traits we used linear regression as implemented in SciKitLearn v1.0 `linear_model.LinearRegression`^8^ and report the coefficient of determination. For discrete-valued traits, we used multinomial logistic regression as implemented in SciKitLearn v1.0 `linear_model.LogisticRegression`^8^. We fit two models: a null model including only intercept or intercept and site, and one including the factor variables. We report the variance explained as the McFadden pseudo-*R*^29^, 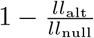, with *ll*_null_ and *ll*_alt_ being the model negative log-likelihood for the null and alternative model respectively.

### 1.5 Calculating relative feature importance

Feature importance in traditional CCA is defined by the correlation of the variables in the reduced space *ρ* = cor(*Y*_1_*f*_1_, *Y*_2_*f*_2_). Unfortunately this notion breaks down in higher dimensions. As we discuss further in the supplemental note, the degree of sharing in MCCA is defined by functions of the cross-correlation matrix in the reduced space,

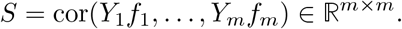

We seek to define an analogous quantity for our graphical model. In MCFA, the data in the reduced (shared) space is given by the posterior mean of *Z*, 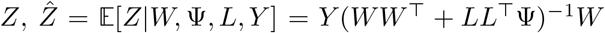. We can also calculate the posterior mean of *Z* conditional on observing a single mode, 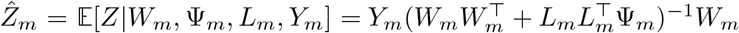. This latter quantity is analogous to the reduced variables *Y*_*m*_*f*_*m*_ in MCCA.

Thus we can summarize the importance of each dimension of the shared space by calculating functions of the cross-correlation of columns of 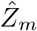,

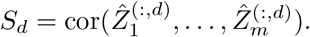

As we show in the supplemental note, the relevant function in our model is the generalized variance |*S*|. The determinant of a correlation matrix is bounded between 0 and 1, with lower values indicating *more* correlation, and higher values *less*. Thus to aid interpretability, we report *ρ*_*d*_ = − log |*S*_*d*_| and re-order columns of *Z* and *W* with decreasing *ρ*_*d*_.

### 1.6 SNP set enrichment analysis

For SNP set enrichment analysis, we broadly follow the approach of CAMERA^10^. In brief, enrichment statistics can be inflated due to correlations in the sample - in this case, linkage disequilibrium between two GWAS SNPs. This results in an under-estimate of the standard error of the enrichment test statistic and an increase in false positives. We calculate the variance inflation factor^10^ by using plink v1.9^11,12^ to estimate linkage disequilibrium between annotation SNPs in 337, 781 unrelated individuals from the UK Biobank^13^. The variance inflation factor is 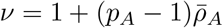, with 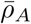 the average person correlation between features in set *A*. We test the known GWAS mean *χ*^2^ statistic 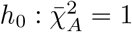 against the alternative 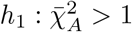. The standard error of the test statistic is 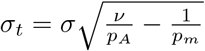 with *σ* the pooled empirical standard deviation of the test statistics.

### 1.7 Preprocessing of the MESA multi-omics pilot dataset

The Multi-Ethnic Study of Atherosclerosis (MESA) is a prospective cohort study with the goal to identify progression of subclinical atherosclerosis^14^. MESA recruited 6,814 participants, ages 45-84 years and free of clinical cardiovascular disease, during 2000-2002. The participants are 53% female, 38% non-Hispanic white, 28% Black, 22% Hispanic and 12% Asian-American. The Multi-Omics pilot dataset includes 30x whole genome sequencing (WGS) through the Trans-Omics for Precision Medicine (TOPMed) Project^15^. Blood samples for multi-omic analysis of participants were collected at two time points (exam 1 and exam 5) and assayed for transcriptomics (RNA-seq in PBMCs, monocytes and T cells), Illumina EPIC methylomics data (whole blood), targeted and untargeted metabolomics data (plasma), and proteomics data (plasma). The MESA Multi-Omics pilot biospecimen collection, molecular phenotype data production and quality control (QC) are described in detail in Aguet et al^16^.

We analyzed individuals from Exam 1 where all five data types were collected and pass QC. All data modalities were inverse rank normalized prior to sample filtering based on the availability of other data types. There were 614 individuals with observations of WGS, RNA-seq, methylation, metabolomics and proteomics that all pass QC. We further removed all features (CpGs, genes, proteins) located on sex-chromosomes, 0-variance features, CpGs with missing data, and CpGs where the probe was within 5 bases of a SNP, leaving us with 6, 042 metabolites, 1, 222 proteins, 19, 034 genes, and 724, 210 CpGs. We analyzed 28 PCs of RNA expression, 39 PCs of methylation, 27 PCs of protein expression and 63 PCs of metabolite, as determined using the aforementioned method. For sample metadata, we leveraged the rich phenotype data available in MESA that were harmonized by the TOPMed Data Coordinating Center^17^. For details on the estimation of sample cell-type proportions from methylation and RNA-seq data, see Kasela et al^18^.

### 1.8 Cross-validation

We used leave-one-out cross-validation (CV) to evaluate our model. The primary reason we chose leave-one-out CV over *k*-fold CV is that our hyperparameter selection method depends on the sample size. With *n* − 1 individuals, the same parameters used for the full inference procedure are likely to be valid. For small *k*, fitting with 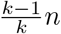 *n* individuals while using the same number of PCs may result in over-fitting in the training set, and using a smaller number of PCs may not capture the same variation as the full model.

To perform cross-validation we hold out a set of individuals, fit the MCFA model, then project the held out individuals into the learned space. If *W*_*tr*_, *L*_*tr*_ and Φ_*tr*_ are the model parameters learned from the training set, the projections of the test data into the learned spaces are given by

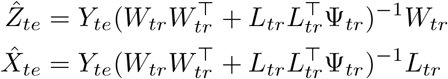

The full data reconstruction is

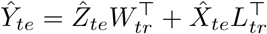

We evaluate model fit by calculating the normalized root mean squared error (NRMSE). In order to provide a fair evaluation across modes with a highly variable number of features, we calculate NRMSE on a per mode basis

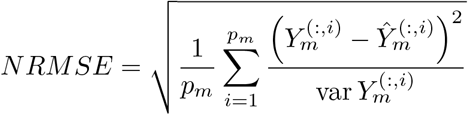

and potential over-fitting can be assessed by comparing the median training set NRMSE against the median test set NRMSE over many cross-validation iterations.

## 2 Acknowledgements

BCB would like to thank Lior Pachter and Nicholas Bray for numerous insightful conversations about CCA over the years. Funding for DAK and BCB is provided by NIA U01AG068880. Funding for BCB is provided by NHGRI K99HG012373 and the Columbia Data Science Institute. Funding for TL and SK is provided by NHLBI R01HL142028. Funding for TL is provided by NIH R01AG057422 and NIMH R01MH106842. TL is a paid adviser or consultant of Variant Bio, GSK, Pfizer and Goldfinch Bio. Whole genome sequencing (WGS) for the Trans-Omics in Precision Medicine (TOPMed) program was supported by the National Heart, Lung and Blood Institute (NHLBI). WGS for “NHLBI TOPMed: Multi-Ethnic Study of Atherosclerosis (MESA)” (phs001416.v1.p1) was performed at the Broad Institute of MIT and Harvard (3U54HG003067-13S1). Centralized read mapping and genotype calling, along with variant quality metrics and filtering were provided by the TOPMed Informatics Research Center (3R01HL-117626-02S1). Phenotype harmonization, data management, sample-identity QC, and general study coordination, were provided by the TOPMed Data Coordinating Center (3R01HL-120393-02S1), and TOPMed MESA Multi-Omics (HHSN2682015000031/HSN26800004). The MESA projects are conducted and supported by the National Heart, Lung, and Blood Institute (NHLBI) in collaboration with MESA investigators. Support for the Multi-Ethnic Study of Atherosclerosis (MESA) projects are conducted and supported by the National Heart, Lung, and Blood Institute (NHLBI) in collaboration with MESA investigators. Support for MESA is provided by contracts 75N92020D00001, HHSN268201500003I, N01-HC-95159, 75N92020D00005, N01-HC-95160, 75N92020D00002, N01-HC-95161, 75N92020D00003, N01-HC-95162, 75N92020D00006, N01-HC-95163, 75N92020D00004, N01-HC-95164, 75N92020D00007, N01-HC-95165, N01-HC-95166, N01-HC-95167, N01-HC-95168, N01-HC-95169, UL1-TR-000040, UL1-TR-001079, UL1-TR-001420, UL1TR001881, DK063491, and R01HL105756. The authors thank the other investigators, the staff, and the participants of the MESA study for their valuable contributions. A full list of participating MESA investigators and institutes can be found at http://www.mesa-nhlbi.org. This publication was developed under a STAR research assistance agreement, No. RD831697 (MESA Air), awarded by the U.S Environmental protection Agency. It has not been formally reviewed by the EPA. The views expressed in this document are solely those of the authors and the EPA does not endorse any products or commercial services mentioned in this publication.

---

**Figure S1:**
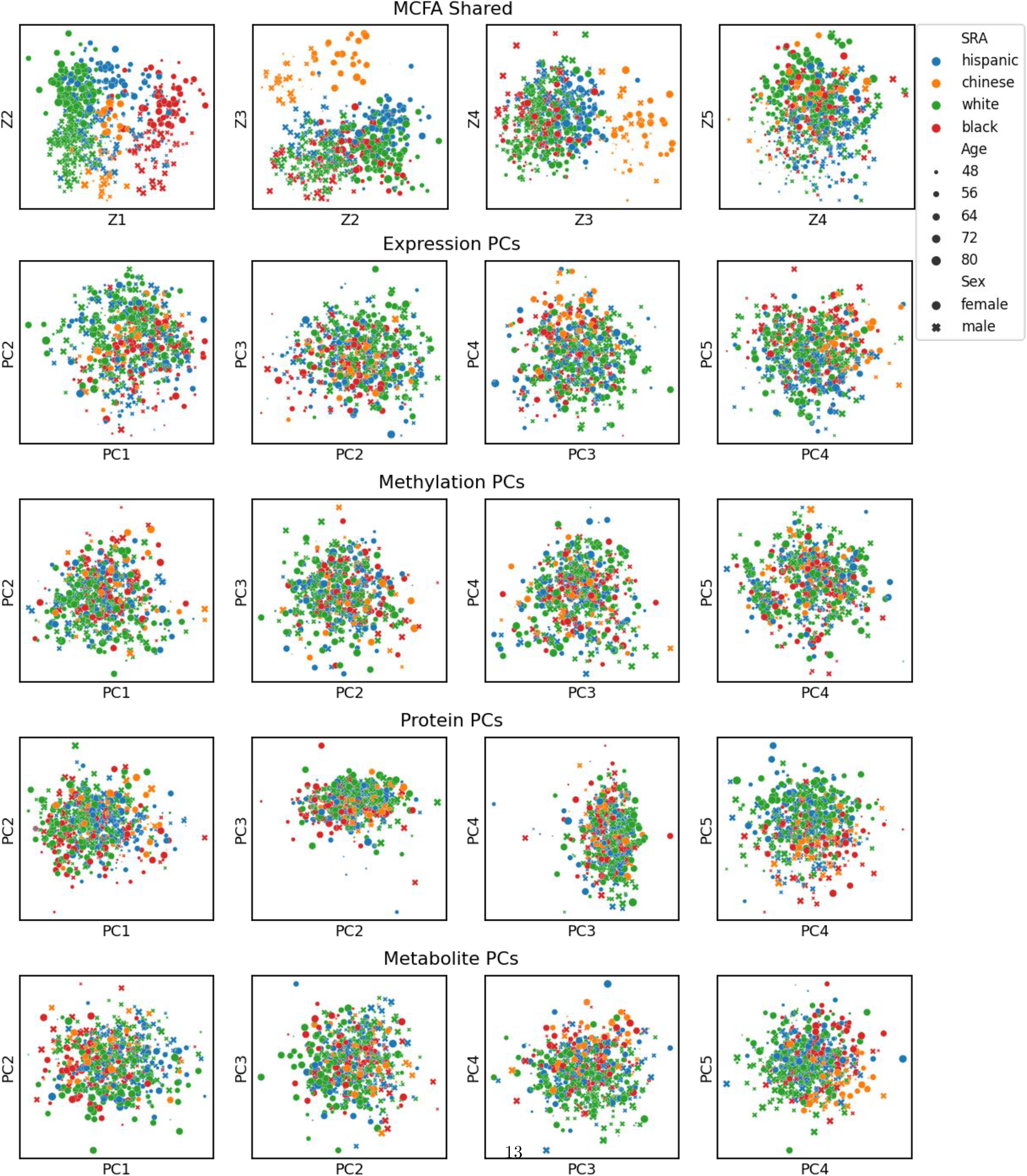
The top shared components learned from the MCFA model clearly reflect self-reported ancestry (SRA), age and sex, while none of the top PCs of any of the datasets show this structure.

**Figure S2:**
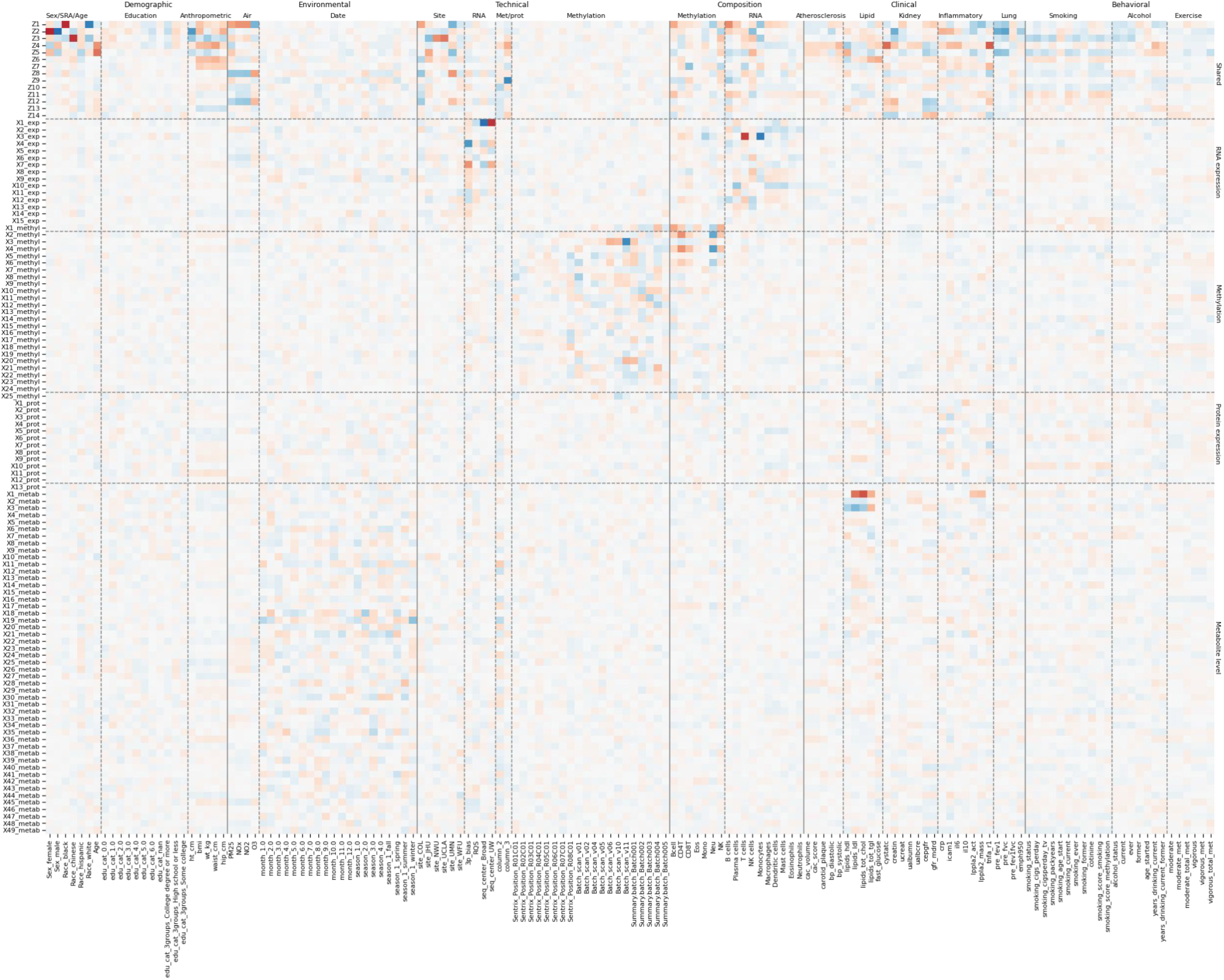
Correlation of each dimension of each learned space (rows) with each metadata factor (columns). Red values indicate positive correlation and blue values negative correlation.

**Figure S3:**
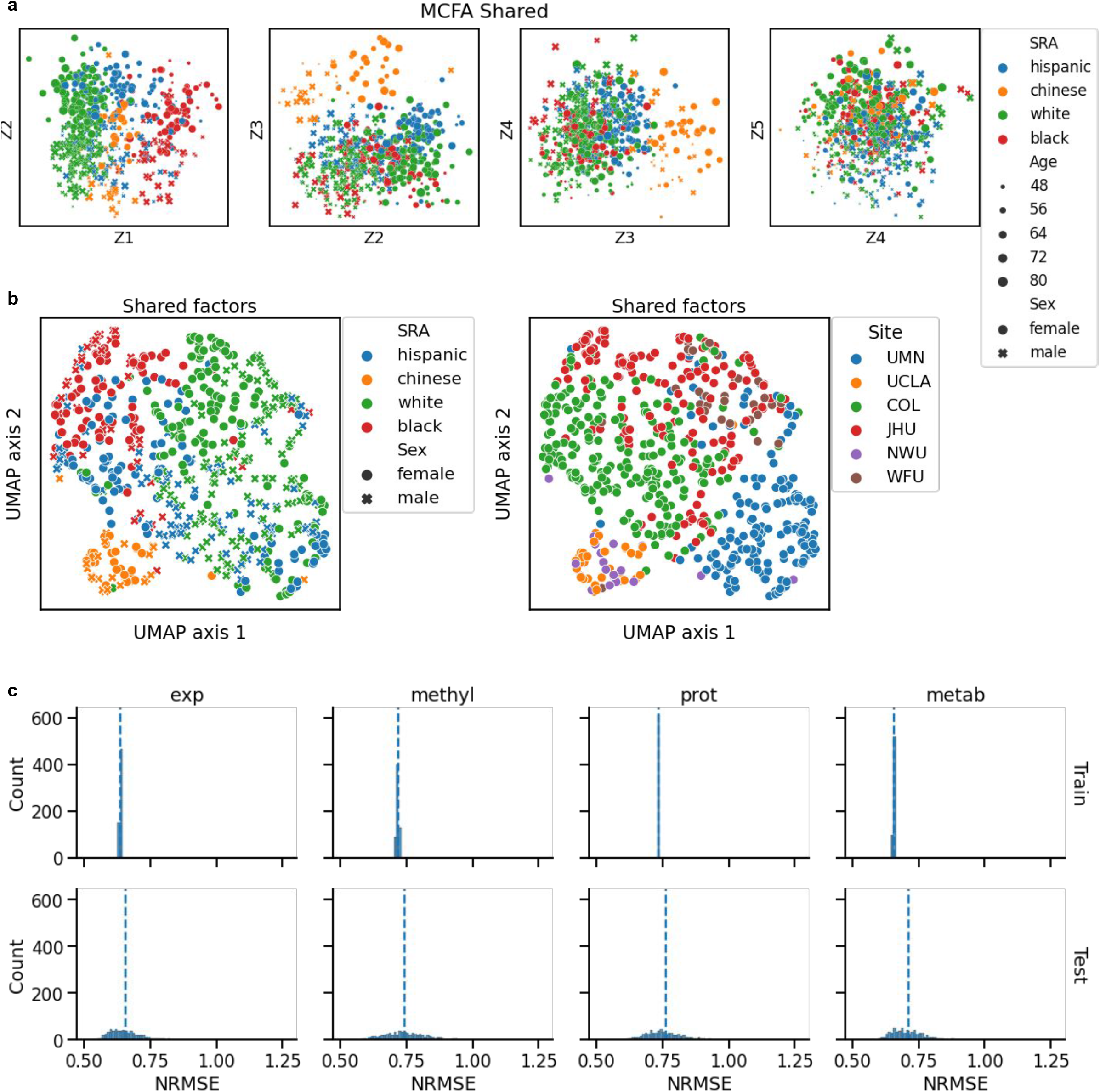
a) The top shared cross-validated MCFA components, annotated by self-reported ancestry (SRA), age and sex. Each point is creating by holding that individual out, fitting MCFA on the remaining individuals, then projecting the held-out individual into the shared space (see Online Methods). b) UMAP embeddings of the cross-validated MCFA components. c) Normalized root mean square error of the 613 training individuals for each dataset (top), versus the held-out individual (bottom), split by data type. Blue dashed line indicates the median.

**Figure S4:**
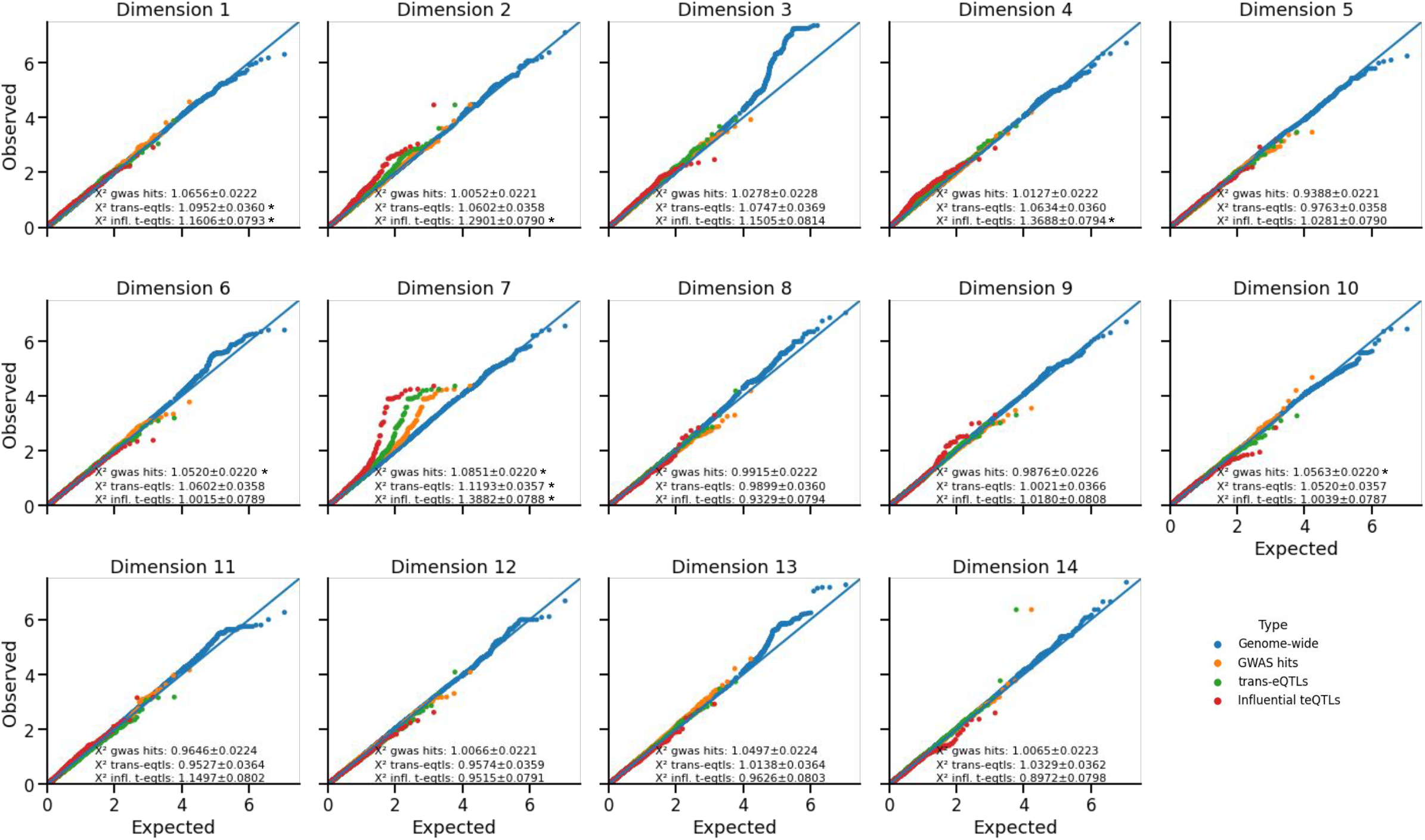
Q-Q plot of GWAS results for each shared factor.

**Figure S5:**
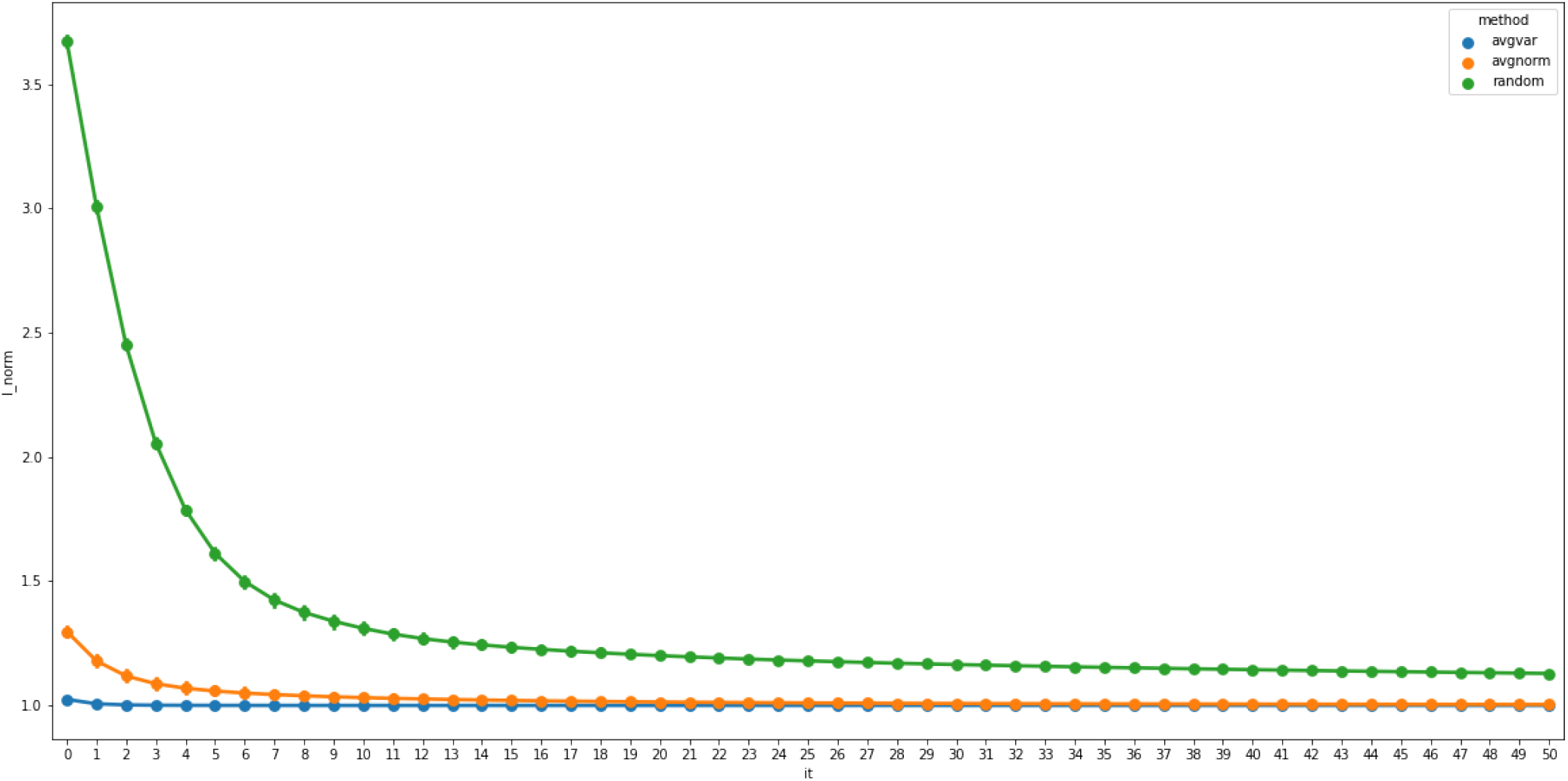
Average normalized model likelihood (y-axis, negative log-likelihood divided by minimum negative log-likelihood) as a function of EM step iteration. We simulated three datasets with *p* = 30, 40, 50 observed features generated by *k*_*m*_ = 8, 11, 15 private and *d* = 10 shared factors and *N* = 1000 individuals in 100 simulations. We compared random, SUMCORR-AVGVAR, and SUMCORR-AVGNORM model initializers. SUMCORR-AVGVAR produced good initial estimates resulting in fast convergence, while other approaches took longer to converge.

## Supplemental Note

### PCA, pPCA and FA

Principal components analysis^1^ is a classic technique for dimensionality reduction. Assume we have *N* samples measured at *p* features each. Let *y*_*n*_ be a *p*-vector denoting the observations for sample *n* and let *Y* = [*y*_1_ : … : *y*_*N*_]^T^ be the corresponding *N* × *p* data matrix. For ease of notation, we assume throughout that each feature has mean 0 but note that this is not a requirement.

There are many ways to derive PCA, but perhaps the most common is to consider the problem of finding a unit projection vector *v*_*i*_ that maximizes the variance in the reduced space *T*_*i*_ = *Y v*_*i*_. The first *k* principal axes *v*_1_, …, *v*_*k*_ are a sequence of orthonormal vectors which successively maximize the variance in the reduced space 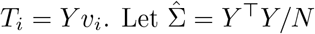 be the empirical covariance matrix of *Y*. PCA solves the following problem:

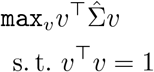

The top-*k* principal axes are thus given by the eigenvectors of 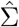 that have the *k* highest eigenvalues. The data in PC space is thus given by the linear projection of the data into this space. Specifically, let *V*_*k*_ = [*v*_1_, …, *v*_*k*_] be the projection matrix, and let *Y* = *U*Λ*V* ^T^ be the singular value decomposition of *Y*. Notice that the eigenvectors of the covariance matrix 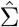 and the right singular values of the data matrix *Y* are the same. The points in PC-space are thus given by *T*_*k*_ = *Y V*_*k*_ = *U*_*k*_Λ_*k*_ where *U*_*k*_ are the the left singular vectors with the *k*-highest singular values.

Tipping and Bishop^2^ introduced a graphical model called probabilistic Principal Components Analysis that provides a generative framework for understanding PCA. The model is as follows:

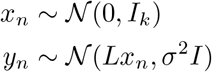

where *L* is a *p* × *k* weight matrix and *σ*^2^ ≥ 0 is the residual noise. They show that the maximum likelihood estimate of the parameters *W* and *σ*^2^ are given by

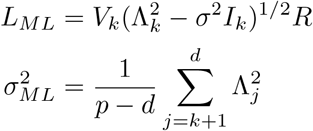

where *R* is an arbitrary *k* × *k* orthogonal rotation matrix. Thus, as *σ*^2^ → 0, *W* represents an orthogonal projection into standard PC space. This defines an equivalence relationship between pPCA and PCA.

Factor analysis is a very similar model, with the only difference being the form of the noise term. Rather than force an isotropic noise model *σ*^2^*I*, factor analysis allows for an arbitrary diagonal positive semi-definite matrix Ψ = diag(*ψ*_1_, …, *ψ*_*p*_) ⪰ 0. This allows each observed feature to have it’s own error variance.

### CCA and pCCA

Now assume each sample is measured on two different sets of conceptually distinct features 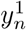 and 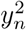 with corresponding *N* × *p*_1_ and *N* × *p*_2_ data matrices *Y*_1_ and *Y*_2_. As before let 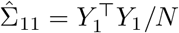 and 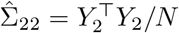 be the empirical covariance matrices for modalities 1 and 2, and let 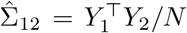 be empirical cross-covariance matrix between the features in each mode. The first set of canonical vectors *f*_1_, *f*_2_ are those that maximize the correlation

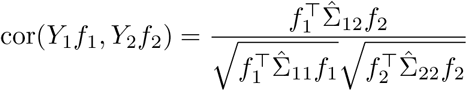

Note that, similarly to PCA, this definition reveals that CCA is a constrained optimization problem:

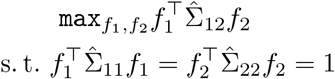

but note that rather than the unit norm constraint *f* ^T^*f* = 1 used in PCA, we have a unit variance constraint 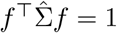. This unit variance constraint allows there to be correlation within the features of a dataset that is not explained by correlation between the features across datasets.

Successive components can be found by projecting out the first canonical component and again maximizing the correlation of the residuals^3^. Equivalently, all components can be found by solving an eigenvalue problem. To see this consider the change of variables 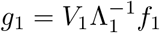 and 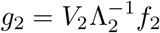. The correlation is now given by:

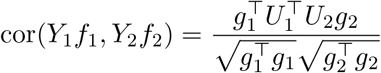

which indicates that *g*_1_, *g*_2_ are the top pair of left-right singular vectors of the matrix 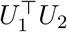. Further components are further singular vectors of 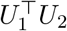, and it’s singular values are the correlations. This also reveals that CCA is equivalent to using PCA to whiten the variables of each data matrix, concatenating them, and then performing PCA again on the whitened, concatenated data matrix.

Likewise to probabilistic PCA, probabilistic CCA is a graphical model that provides a generative framework for thinking about CCA^4^. The model is as follows:

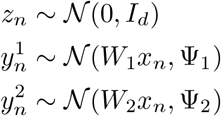

similarly to FA and pPCA, we sample a *d*-dimensional random normal hidden vector, pass it through a weight matrix, and add random noise. We have two weight matrices *W*_1_ and *W*_2_ of shape *p*_1_ × *d* and *p*_2_ × *d*, and two noise matrices Ψ_1_ and Ψ_2_, however in this case these noise matrices are arbitrary positive semi-definite matrices (Ψ_•_⪰ 0). Bach and Jordan show that the maximum likelihood estimate of the parameters of pCCA can be determined from the CCA solution:

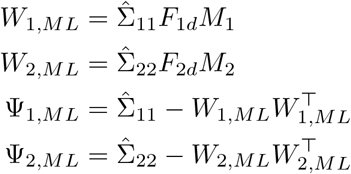

where *F*_•*d*_ = [*f*_•1_; … ; *f*_•*d*_] are the first *d* canonical directions and *M*_1_, *M*_2_ are arbitrary matrices with spectral norm less than 1 such that *M*_1_*M*_2_ = *ρ*_*d*_.

### Multi-set canonical correlation analysis

Now rather than having two sets of conceptually distinct features for each sample, assume we have *M* different conceptually distinct sets of features 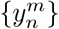 with corresponding *N* × *p*_*m*_ data matrices {*Y*_*m*_}. In MCCA, we are still interested in finding projection vectors {*f*_*m*_} which map our high dimensional data into a one-dimensional space, however there are many formulations that are equivalent to classical CCA with two datasets. Let 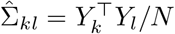 be the empirical cross-covariance matrix between the features in dataset *k* and dataset *l*. The covariance of the data in the reduced space is given by

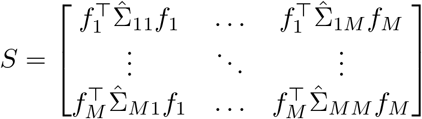

The various formulations of MCCA correspond to optimizing different objective functions *J* (*S*) subject to the a constraint function 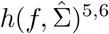. In brief, possible objective functions include:

- SUMCOR: Maximize the sum of pairwise correlations: 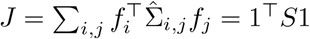
- SUMSQCOR: Maximize the sum of squares of pairwise correlations: 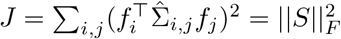
- MAVAR: Maximize the largest eigenvalue of S: *J* = *λ*_1_(*S*)
- MINVAR: Minimize the smallest eigenvalue of S: *J* = *λ*_*d*_(*S*)
- GENVAR: Minimize the determinant of S, also known as the generalized variance: *J* = |*S*| = ∏_*i*_ *λ*_*i*_(*S*)

while possible constraints include:

- VAR: The canonical directions each have unit variance: 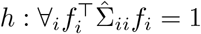
- AVGVAR: The canonical directions have unit variance on average: 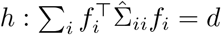
- NORM: The canonical directions each have unit norm: 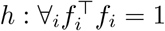
- AVGNORM: The canonical directions have unit norm on average: 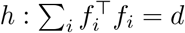

It is straightforward to see that any of the 5 listed objective functions could be combined with either of the first two constraints to create 10 optimization problems that are equivalent to CCA in the two-dataset case. The final two constraints correspond to relaxations of the unit variance constraint which can reveal a simpler optimization problem in some cases, and which does not suffer from being trivially satisfiable when *p > N*.

Some of these can be fit by solving eigenvalue problems, while others require more complicated iterative methods. Of particular note are the SUMCOR and GENVAR objectives. The GENVAR objective was the first considered MCCA approach^7^, where a simple solution for the *M* = 3 and *p*_1_ = *p*_2_ = *p*_3_ = 2 case is given. GENVAR is a particularly natural way of thinking about MCCA - it is a single value that represents the multidimesnioal scatter of points in space^8^. Smaller values of the generalized variance indicate less scatter, and thus higher “correlation” of the points in the reduced space. Despite this, is has received relatively little attention as a method for MCCA, perhaps because it is challenging to fit^6^. On the other hand, most attention has been focused on the SUMCOR objective^9^, which can be solved easily with the AVGVAR and AVGNORM constraints. SUMCOR with AVGVAR constraint can be solved via a simple two-stage procedure: first whiten each data matrix, concatenate the whitened features, and then perform PCA on the whitened, concatenated features. SUMCOR with AVGNORM constraint is even simpler to solve: simply perform PCA on the concatenated feature set. In the two dataset case, this latter method is sometimes called “diagonal CCA” and forms the basis of the original integration approach used in Seurat^10^ as well as many sparse CCA approaches^11^. This is also closely related to group factor analysis approaches for multi-modal data, see for example^12^ and references therein. The equivalence to PCA on the concatenated feature set makes it straightforward to see that the NORM-based constraints involve an implicit assumption that shared factors are responsible for both covariation across features in different modes and between features within a mode.

### Probabilistic graphical model for multi-set CCA

Here, we describe a probabilistic graphical model for multi-set CCA (pMCCA). Note that while this model is an option in our software package, we derive and discuss it primarily to draw connection to traditional multiset CCA. The full model which includes simultaneous factor analysis of the private spaces is described in the next section. Unlike traditional CCA, pCCA has one single obvious generalization to multiple datasets:

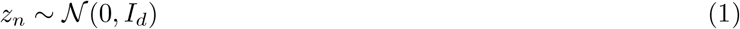

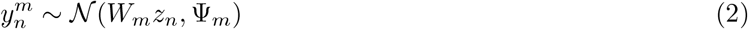

where again we have weight matrices *W*_*m*_ of shape *p*_*m*_ × *d* and arbitrary positive semi-definite noise matrices Ψ_*m*_ ⪰ 0. This model is illustrated as a plate diagram in Figure S6.

We will now see that pMCCA is related to MCCA in the following ways:

**Figure S6:**
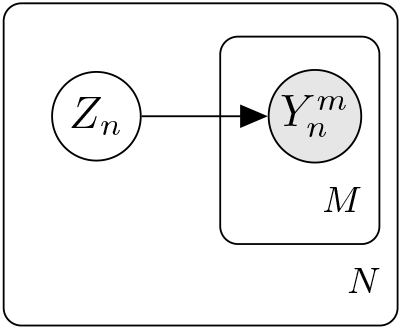
Probabilistic multi-set canonical correlation analysis (pMCCA). For each individual *n* (the outer plate), we sample *z*_*n*_ from the *d*-dimensional unit Gaussian, *z*_*n*_ ∼ 𝒩 (0, *I*_*d*_). For each dataset *m* (the inner plate) we sample the features of that dataset from a *p*_*m*_-dimensional unit Gaussian, 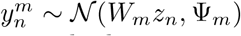. Here 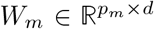 is the transformation weight matrix and 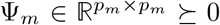 is the residual covariance matrix.

#### Observation 1.

*The maximum likelihood solution to the pMCCA model corresponds to the GENVAR objective with VAR constraint in the M* = 3 *dataset case*.

#### Observation 2.

*The maximum likelihood solution to the pMCCA model does not correspond to any of the listed MCCA formulations in the M* ≥ 4 *case*.

Let 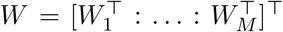 sbe the stacked weight matrices and Ψ = diag(Ψ_1_, …, Ψ_*M*_) be the block diagonal covariance matrix. The model covariance is given by:

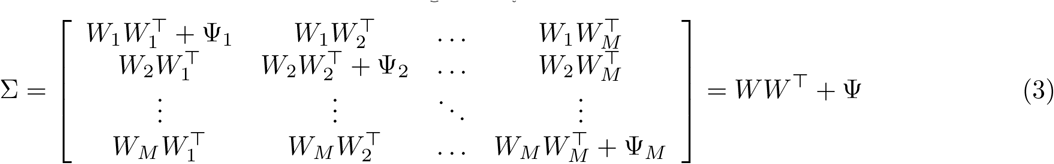

Let *Y* = [*Y*_1_ : … : *Y*_*M*_] so that the empirical covariance matrix can be written 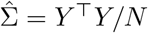. The model negative log-likelihood is given by:

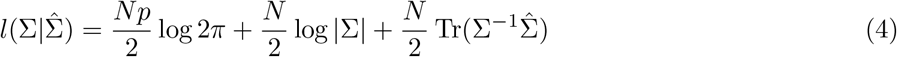

where *p* = ∑_*m*_ *p*_*m*_ is the total number of features from all datasets. It is straightforward to see that many arguments from Bach and Jordan^4^ carry over to the multi-set case, thus we refer readers there for proofs. In particular we have

#### Lemma 1.

*At a stationary point of the likelihood* 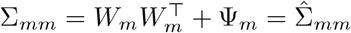.

Thus, at a stationary point

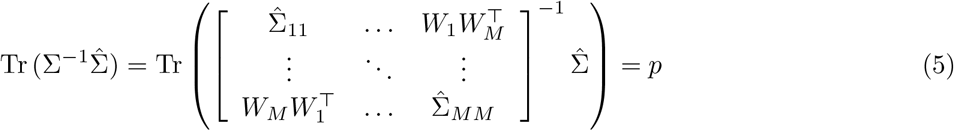

so that the models minimum negative log-likelihood is proportional to the log generalized variance of the model:

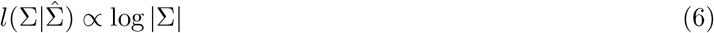

Moreover, Lemma 1 allows us to further factorize Σ

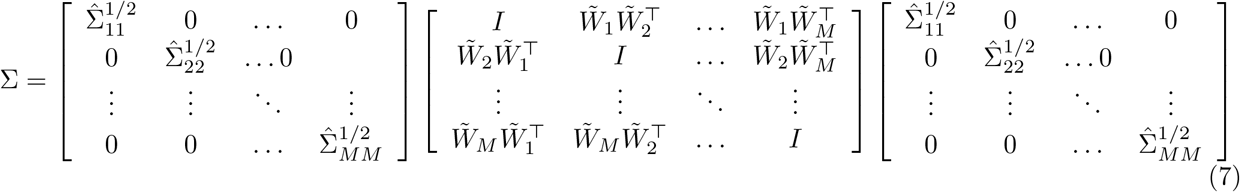

and so

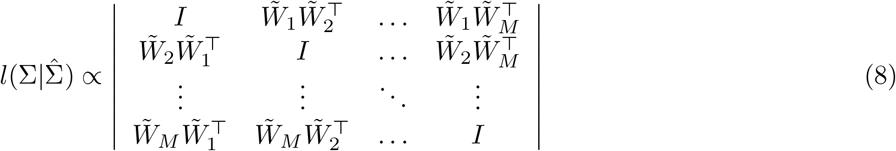

where 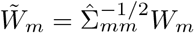. Notice that the off-diagonal blocks are the cross-covariance matrices in the model of individually-whitened datasets. If there exists a set of projection vectors *f*_*m*_ such that 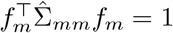 and the projection of the data into the space spanned by *f*_*m*_ has covariance equal to the above, then minimizing the above determinant solves the GENVAR MCCA objective with VAR constraint. Let 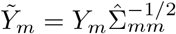 be the whitened datasets and let 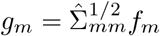 be the change of variables that gives 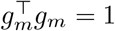. The projection in MCCA space is given by 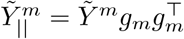. We seek *g*_*m*_ such that

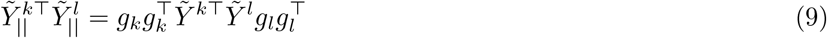

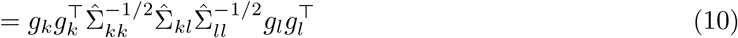

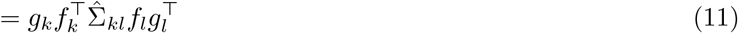

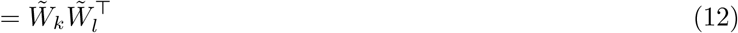

Thus we must satisfy

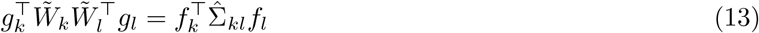

Notice that for *d* = 1, the left and right side are scalars. This means we can express our necessary criterion as

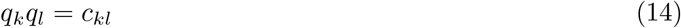

For *M* = 3, this results in a system of 3 equations in 3 unknowns, which has solutions of the form 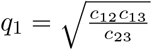. Note that these can be found by setting 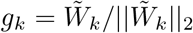 which yields 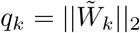, For *M* > 3, there are more equations than unknowns and they cannot be mutually satisfied in general. Note also that a similar argument can be used in the *d* > 1 case to show that this system is not satisfiable even for *M* = 3. Thus, unlike CCA and pCCA, fitting *d* > 1 components is not equivalent to iteratively fitting single components and projecting them out.

### Multiset Correlation and Factor Analysis

The residual covariance matrices Ψ_*d*_ deserve additional attention. Put simply, these matrices represent the residual structure in each modality after accounting for shared structure across modes. Instead of allowing this matrix to be arbitrary, we can instead think of this matrix as having some additional structure. For example, Ψ_*d*_ might be the sum of a low rank and an isotropic covariance matrix. This suggests that we add an additional latent variable to each dataset, corresponding to a factor model for the “private” structure (e.g. not shared with other datasets). Specifically, we modify pMCCA such that for each dataset, we additionally sample a latent variable from a *k*_*m*_-dimensional unit Gaussian. The observed data are then sampled from a multi-variate Gaussian where the mean is a linear combination of both private variables, but the residual covariance matrix is now diagonal.

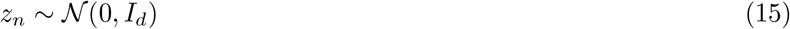

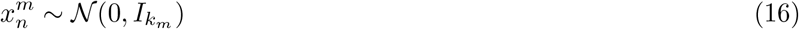

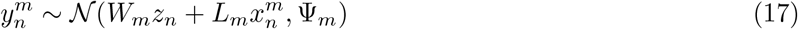

where *L*_*m*_ are the *k*_*m*_ × *p*_*m*_ private space loading matrices and 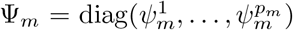 are the diagonal residual covariance matrices. Note that in general we allow the entries on the diagonal of Ψ_*m*_ to take different values, similarly to factor analysis. One could additionally constrain the entries of the diagonal to be the same, Ψ = *σ*^2^*I*_*p*_, similar to pPCA.

**Figure S7:**
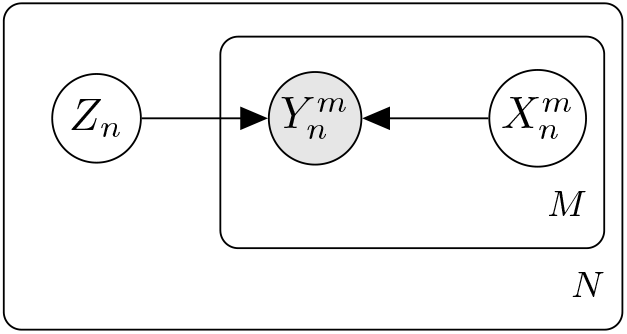
Multset correlation and factor analysis as a plate diagram. For each individual *n* (the outer plate), we sample *z*_*n*_ from the *d*-dimensional unit Gaussian, *z*_*n*_ ∼ 𝒩 (0, *I*_*d*_). For each mode *m* (the inner plate) and individual *n* we sample 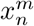 from a *k*_*m*_-dimensional unit Gaussian. The observed features of that mode are then sampled from a *p*_*m*_-dimensional unit Gaussian, 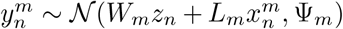. Here 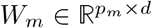 is the transformation weight matrix, *L*_*m*_ is the private-space loadings matrix, and 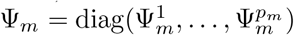 is the diagonal residual covariance matrix.

This model can be fit via a straightforward application of the expectation-maximization (EM) algorithm^13^. We first derive conditional expectation of the log-likelihood, ℒ for the model under the generative process specified in Figure S7. For convenience, let

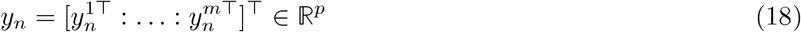

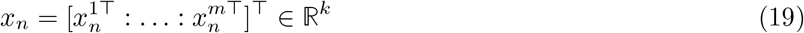

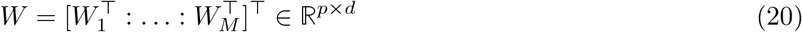

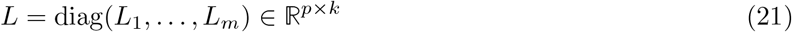

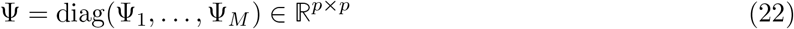

where *k* = ∑_*m*_ *k*_*m*_. At a given time step *t* during the computation of the EM algorithm, let the conditional expectation for given latent variables *z*_*i*_ and *x*_*i*_ be 𝔼[·|*W*_*t*_, Ψ_*t*_, *L*_*t*_, *y*_*i*_] = ⟨·⟩.

The conditional expectation of the log-likelihood (E-step) is:

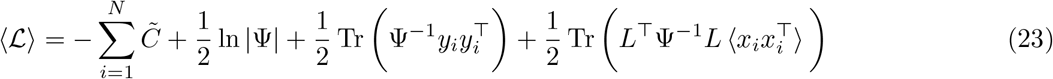

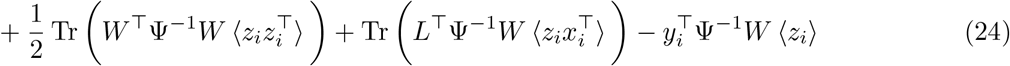

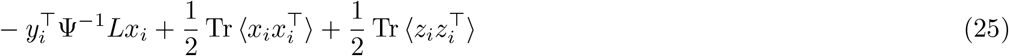

At a given timestep *t*, we compute the update of parameters *t* + 1 by differentiating ℒ with respect to *W*_*t*_, *L*_*t*_, and Ψ_*t*_, and setting the derivative of the corresponding expected log-likelihood to 0. The following update steps are derived using standard matrix differentiation results^14^.

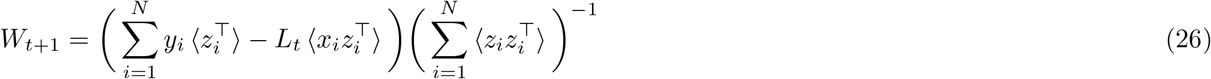

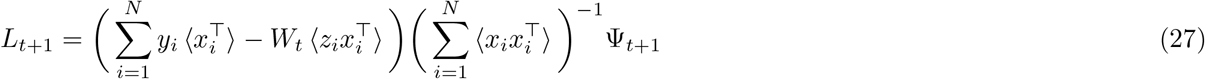

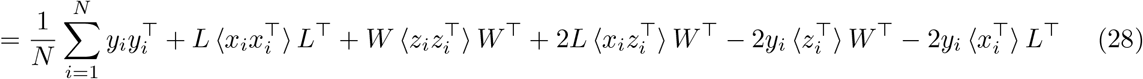

